# Marginal Zone B Cells are Necessary for the Formation of Anti-Donor IgG After Allogeneic Sensitization

**DOI:** 10.1101/2022.09.23.509210

**Authors:** Melissa A. Kallarakal, Gregory Cohen, Francis I. Ibukun, Scott M. Krummey

## Abstract

The formation of anti-MHC antibody is a significant barrier to improved outcomes in organ transplantation. Patients with pre-formed anti-HLA antibodies have limited options for suitable donors, and the formation of donor-specific anti-HLA antibodies after transplantation is a harbinger of graft rejection. Despite the recognized importance of anti-HLA antibodies, the mechanisms responsible for the differentiation of B cells after exposure to allogeneic antigen are poorly understood. In order to evaluate the differentiation of B cells in response to allogeneic antigen, we used a model of H-2^b^ C57/Bl6 sensitization with H-2^d^ antigen. We found that although the formation of anti-H-2^d^ IgG was robust, few class switched B cells and germinal center B cells were formed. Sensitization induced weak expression of classical memory B cell markers, but we observed populations of CD21^+^ and IRF4^+^ B cells, that corresponded to an increase in the frequency of marginal zone phenotype B cells after sensitization. Depletion of marginal zone B cells prior to sensitization resulted in a significant dimunition of anti-H-2^d^ IgG and also fewer germinal center B cells. These results demonstrate a previously unappreciated role for marginal zone B cells as a reservoir of alloreactive B cells that are activated by allogeneic antigen.

## INTRODUCTION

Solid organ transplantation is curative therapy for end-stage organ failure due to a wide range of diseases (1,2). However, the successful impact of of transplantation is limited by the formation of anti-HLA antibodies, termed HLA sensitization (3,4). For patients awaiting transplantation, HLA sensitization can dramatically limit the number of compatible donors, and the formation of anti-donor HLA antibodies after transplantation leads to graft rejection and loss (5,6). Thus, further mechanistic insight into the cellular and molecular pathways that lead to HLA sensitization could lead to improved risk stratification of patients prior to transplant and pathway-specific treatments for graft rejection (7–10).

Although plasma cells are major source of DSA, the pathways by which B cells differentiate into plasma cells after encountering allogeneic antigen are poorly understood. Naïve mature B cells differentiate into two distinct subets, follicular (FO) and marginal zone (MZ) populations. Relative to FO B cells, which are IgD^+^ and reside within the follicle of secondary lymphoid organs, MZ B cells are IgM^+^ cells that reside along the follicule periphery in the marginal zone (11,12). MZ B cells are described as innate-like, owing to their capacity to rapidly respond to bloodborne pathogens and pattern recognition receptors (PRR) such as the TLR4 agonsit lipopolysaccharide (LPS) (13–15). Relative to FO B cells, MZ B cells are poised to rapidly form IgG secreting plasma cells (15,16). Despite these differences, there is significant functional overlap between FO and MZ B cells, as each can acquire the opposite phenotype in certain contexts, form germinal center (GC) B cells and memory B cells.

In transplantation, there is growing evidence that innate-like and MZ B cells play a role in HLA sensitization and graft rejection. Depletion of MZ B cells prolongs heart graft survival after T cell depletion, and marginal zone B cells are required for antibody responses against a red blood cell alloantigen (17–19). Recent studies found that in rejecting hearts and kidneys, infiltrating B cells populations are responsive to innate stimuli and have self-reactivity.

In this study, we sought to understand which B cell populations are important for the formation of allogeneic IgG post-sensitization. In this study, we sought to gain deeper understanding of the mechanisms of HLA sensitization by tracking antigen-specific B cells after exposure to allogeneic antigen. We found a critical role for MZ B cells in the formation of antidonor IgG and plasma cells.

## METHODS

### Mice and Allogeneic Sensitization

C57Bl/6 (H-2^b^) and Balb/c (H-2^d^) mice were obtained from Jackson Laboratories or bred in-house. The study was conducted in accordance with the reocmmendations in the Guide for the Care and Use of Laboratory Animals. The protocol was approved by the Animal Care and Use Committee at Johns Hopkins University. Animals were housed in specific pathogen-free animals facilities. For allogeneic sensitization, Balb/c H-2^d^ splenocytes were harvested and processed to a single-cell suspension. Approximately 20×10^6^ cells were administered via intraperitoneal (IP) injection into C57Bl/6 H-2^b^ mice. This dosage was repeated once per day for 3 days to give ample antigen exposure.

### MHC Tetramer Enrichment and Flow Cytometry

Biotinylated MHC monomers with human β2-microglobulin specific for H-2L^d^ β-galactosidase (TPHPARIGL) were obtained from the NIH Tetramer Core Facility and tetramerized with streptavidin-PE (Prozyme). Decoy reagent was made according to published protocols (20). Briefly, streptavidin-PE (Prozyme) was conjugated to AlexaFluor647 using the Antibody Labeling Kit (ThermoFisher). The concentration decoy reagent was quantified using a NanoDrop. For flow cytometry and magnetic enrichment were performed using FACS buffer comprised of 1X PBS (without Ca^2+^/Mg^2+^, pH 7.2), 0.25% BSA, 2 mM EDTA, and 0.09% azide. Single cell suspensions of splenocytes were incubated with decoy reagent, followed by MHC tetramer, and enriched using anti-PE beads and paramagnetic LS columns (Miltenyi) according to published protocols. The column bound fraction was analyzed as the MHC tetramer enriched fraction, and the flow-through was collected as the unbound or bulk fraction.

### Flow Cytometry Analysis

B cell populations were identified as live (Zombie Near IR^-^; Biolegend), lineage (Thy1.2, NK1.1, F4/80)^-^, and CD19^+^B220^+^. Both fractions were stained with surface antibodies for 30 min at room temperature and intracellular antigens were assessed using the Transcription Factor Staining Kit (eBiosciences). Populations were identified using antibodies against CD19, CD21/35, CD23, CD38, CD45R (B220), CD73, CD80, F4/80, GL7, IgD, IgM, IRF4, NK1.1, PD-L2, and/or Thy1.2 (Biolegend, ThermoFisher, BD Biosciences). All experiments were acquired using a 4L Cytek Aurora, and gating was performed using FlowJo (BD Biosciences). For dimensionality reduction analysis, manually-gated bound H-2L^d^ tetramer^+^ IgD^lo^ B cell events were imported into R (4.1.1) through CytoML (2.40), flowWorkspace (4.4.0), and flowCore (2.4.0). The data were further analyzed using CATALYST (1.16.2) with FlowSOM (2.0.0) clustering and ConsensusClusterPlus (1.56.0) meta clustering. UMAP dimensionality reductions were generated with scatter (1.20.1). Visualizations were generated in ggplot2 (3.3.5) and ComplexHeatmap (2.8.0).

### Cellular H-2^d^ Crossmatch

Serum was collected from C57Bl/6 mice before or at indicated times after allogeneic sensitization. Single cell suspensions of Balb/c splenocytes were prepared, and 1×10^6^ splenocytes were incubated with 1 μL of serum for 1 h at room temperature. Cells were washed 2X with FACS buffer, followed by anti-mouse IgM, anti-mouse IgG, and anti-B220.

### Depletion of Marginal Zone Subsets

Depletion of the Marginal Zone population in vivo was done by treating with 100 μg each anti-CD11a/anti-CD49d or IgG isotype control (BioXCell) on days −4 and −2 prior to sensitization.

### Statistical Analysis

All data points represent individual animals. Expression levels were compared using Student’s t-test (two-tailed) or ANOVA where appropriate. Statistics were performed using GraphPad Prism 9. Significance was determined as *p<0.05, **p<0.01, ***p<0.001, ****p<0.0001.

## RESULTS

### Allogeneic sensitization of H-2^d^ cells induces anti-donor antibodies and modest frequencies in isotype-switched B cells

In order to assess the dynamic changes in the B cell compartment after allogeneic sensitization, we sensitized C57Bl/6 (H-2^b^) hosts are sensitized with allogeneic H-2^d^ Balb/c antigen (Figure 1A). This model allows the assessment of anti-H-2^d^ IgM and IgG via cellular crossmatch and antigen-specific B cells using a MHC Class I H-2L^d^ tetramers. We assessed the formation of anti-donor IgM and IgG after sensitization. We found that anti-donor IgM was significantly increased at day 7 and diminished at day 14 and 21 (Figure 1B). Anti-donor IgG increased at day 7 and remained elevated at day 14 and 21 (Figure 1C). Thus, exposure to allogeneic antigen induces anti-donor IgM and IgG responses.

**Figure 1.**
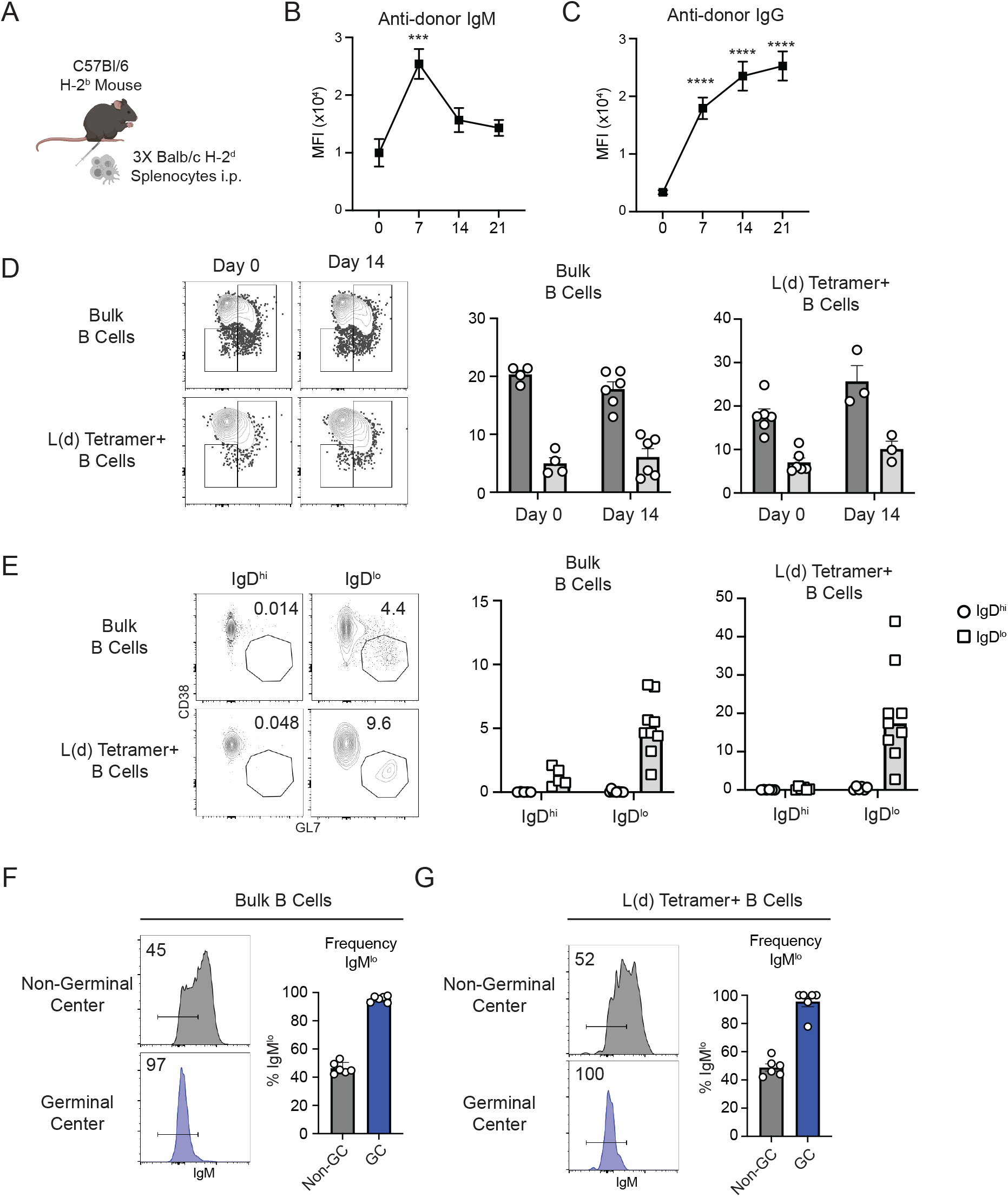
Allogeneic sensitization induces IgG and IgM antibodies and modest germinal center populations. (A) C57Bl/6 mice were sensitized by intraperitoneal administration of Balb/c (H-2^d^) splenocytes (days 1,2, and 3). (B) Anti-donor IgM and (C) anti-donor IgG were assessed in the serum on day 0-21 post-transplant. (D) The expression of IgM and IgD on bulk CD19^+^ and H-2L^d^ tetramer^+^ B cells on day 0 and 14 post-sensitization. (E) The frequency of CD38^lo^GL7^+^ germinal center B cells on day 0 and 14 post-sensitization. The frequency of IgMlo cells among germinal center B cells and non-germinal center (IgD^lo^CD38^+^GL7^-^) (F) bulk CD19+ B cells and (G) H-2L^d^ tetramer^+^ B cells. Each data point represents an individual mouse, and all summary data represents compiled data from 2-3 independent experiements. ***p<0.001, ****p<0.0001.

We next assessed the isotype class switching of total and H-2L^d^-specific B cells. We found no significant changes in the isotype profile of B cells on day 14 relative to day 0 (Figure 1D). We also assessed the formation of germinal center CD38^lo^GL7^+^ B cells, and found that within the IgD^lo^ compartment, a modest frequency (10-20%) of germinal center B cells were formed (Figure 1E). Although the overall frequency of isotype class switching was not different (Figure 1D), the germinal center B cells were universally IgM^lo^ Thus, sensitization induces a durable donor-specific IgG formation, but relatively modest number of class switched B cells.

### Allogeneic sensitization induces few classical memory B cell populations

Antigen-experienced B cells are typically defined by the expression of three markers, CD73, PD-L2 and CD80. We assessed the expression of these receptors in bulk and antigenspecific H-2L^d^ B cells on day 21 after allogeneic sensitization by comparing naive IgD^hi^ and activated IgD^lo^ populations. Although CD73 and PD-L2 were modestly increased in bulk IgD^lo^ B cells relative to IgD^hi^ B cells (Figure 2A), there was no difference in the expression of these receptors in antigen-specific B cells (Figure 2B). Thus, although these receptors can be used to distinguish antigen-experienced and memory B cells induced after infection or hapten immunization, they do not appear to be specifically upregulated after allogeneic sensitization.

**Figure 2.**
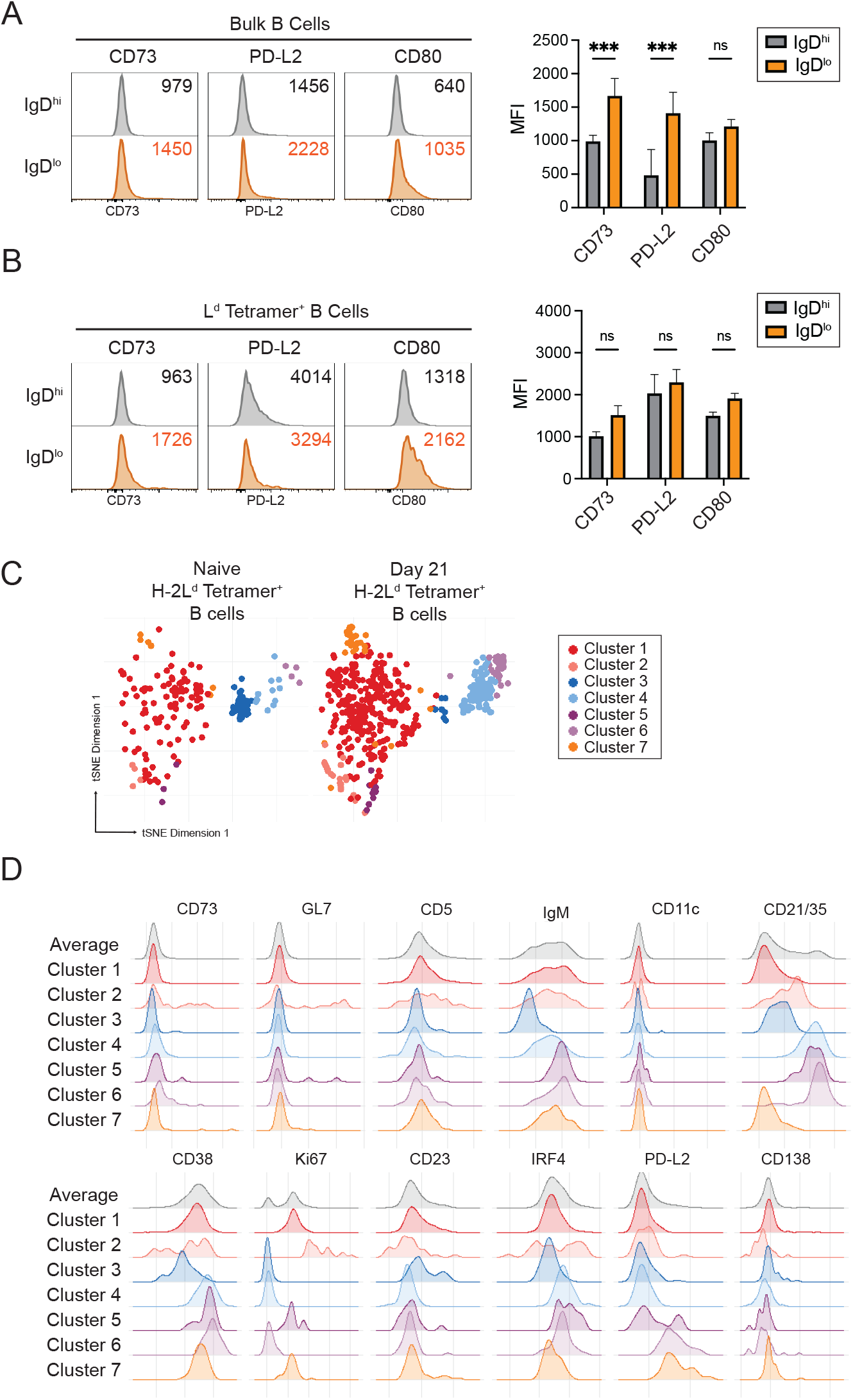
Dimensionality reduction analyses reveals B cell populations in sensitized mice. (A-B) The expression of anti-experienced B cell markers CD73, CD80, and PD-L2 were assessed on (B) bulk CD19+ B cells and (B) H-2Ld tetramer+ B cells. (C) Phenotypic profiles of naïve and day 21 H-2Ld tetramer+ B cells were clustered and visualized using t-SNE. (D) Histograms of phenotypic markers in each individual cluster. Each data point represents an individual mouse, and all summary data represents compiled data from 2-3 independent experiements. ***p<0.001.

In order to evaluate the phenotypic changes that occur after allogeneic sensitization, we evaluated the phenotype of antigen-specific B cells on day 0 and day 21 after sensitization using clustering algorithms and visualization with t-SNE. We found that x clusters appeared to be enriched in day 21 versus day 0. We found that among seven clusters, three expressed very high levels of CD21/35. These clusters were also relatively high for IgM, low expression of CD23 and Ki67, and expressed high/intermediate levels of IRF4. Thus, this unbiased data provided a potential phenotype of antigen-experienced B cells after allogeneic sensitization.

### Allogeneic sensitization induces an increased frequency of marginal zone phenotype B cells that have high IRF4 expression

We next evaluated whether the phenotype of enriched clusters corresponded to populatiosn changes detected by manual gating. The complement receptor CD21 (CR2) is a marker of marginal-zone phenotype B cells, so we evaluated the frequency of CD21+ B cells among IgD^lo^ antigen-specific and bulk B cells. We found that the frequency of marginal-zone B cells increased at day 21 and 42 post-sensitzaiton in both subsets (Figure 3A). We also evaluated the expression of the transcription factor IRF4, which is both implicated in the development of marginal zone B cells and reflects the antigen-strenght of stimulation of lymphocytes. We found that IRF4 expression was signiflicantly enriched in the CD21 ^+^ bulk and antigen-specific B cells.

**Figure 3.**
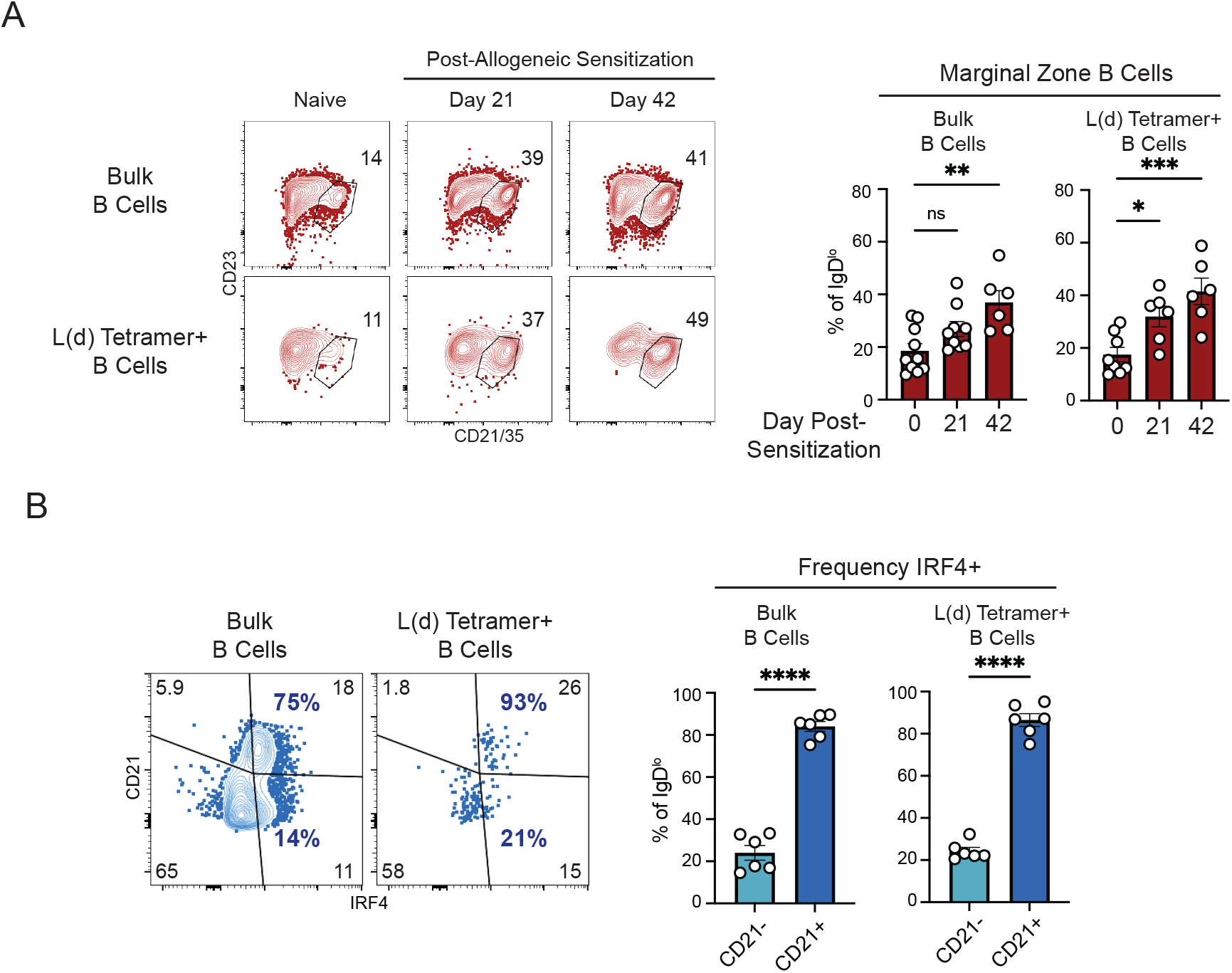
The frequency of CD21+IRF4+ marginal zone B cells increases after allogeneic sensitization. (A) The frequency of CD21+ bulk CD19+ and H-2Ld tetramer+ B cells on day 0, 21, and 42 post-transplant. (B) The freuqencey of IRF4+ B cells among CD21+ and CD21-bulk CD19+ and H-2Ld tetramer+ B cells. Each data point represents an individual mouse, and all summary data represents compiled data from 2-3 independent experiements. ***p<0.001, ****p<0.0001.

### Depletion of the marginal zone B cells significantly restrains the formation of anti-donor IgG and germinal center B Cells

In order to evluate whether the marginal zone B cells are a precursor population of allogeneic B cells or differentiate over time, we assessed the formation of anti-donor antibody in wild-type or marginal zone B cell depleted C57Bl/6 hosts (Figure 4A-B). We found that at day 7, the anti-donor IgM was diminished in MZ-depleted hosts (Figure 4C). Anti-donor IgG was diminished in MZ-depleted hosts at both day 7 and Day 14 (Figure 4D). Thus, it appears that the marginal zone B cell population is a precursor population for differentiating allogeneic B cells.

**Figure 4.**
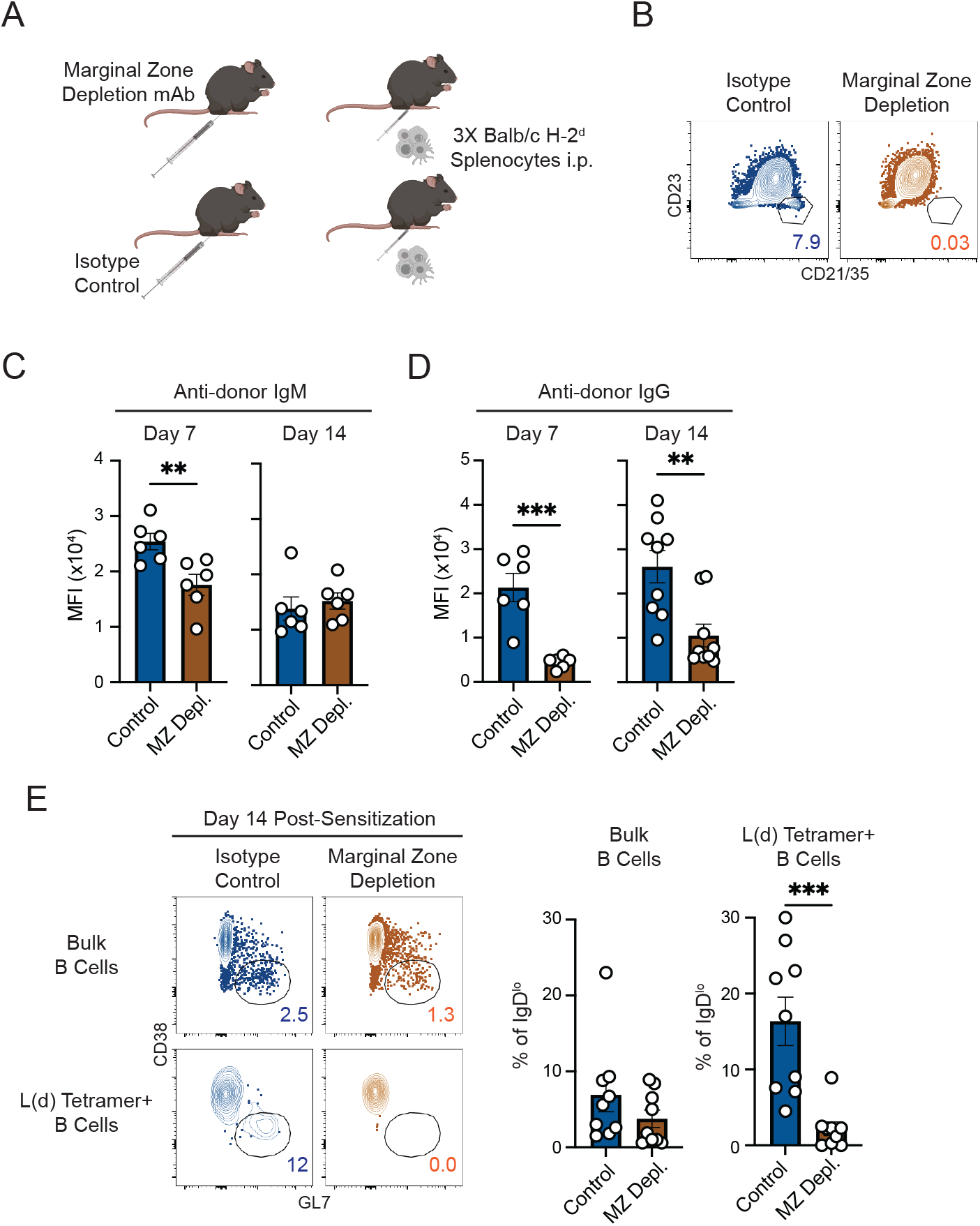
Depletion of marginal zone B cells diminishes anti-donor IgG and the formation of germinal center B cells. (A) Mice were treated with marginal zone depletion antibodies or isotype control prior to sensitization. (B) Representative flow plots of CD21+ marginal zone B cells after treatments. (C) Anti-donor IgM in the serum on day 7 and 14 in mice marginal zone depleted (MZ Depl.) or isotype control (Control) mice. (D) Anti-donor IgG in the serum on day 7 and 14 in mice marginal zone depleted (MZ Depl.) or isotype control (Control) mice. (E) The freuqencey of germinal center B cells among bulk CD19+ and H-2Ld tetramer^+^ B cells on day 14 post-sensitization. Each data point represents an individual mouse, and all summary data represents compiled data from 2-3 independent experiements. **p<0.01, ***p<0.001.

We next assessed the phenotype of bulk and antigen-specific B cells in wild-type and marginal zone depleted hosts. We found that the frequency of germinal-center B cells was slightly but not statistically diminished in bulk B cell ppoulations, the formation of germinal center B cells was significantly diminished in antigen-specific B cells. These data suggest that the majority of the germinal center B cells, along with class switched plasmablasts and plasma cells, originate from the marginal zone B cell compartment.

## Discussion

HLA sensitization remains a significant barrier to improved outcomes for transplant patients prior to and after transplantation. Deeper understanding of the mechanisms by which HLA sensitization occurs could lead to better risk stratification of transplant patients and original approaches to prevent rejection. We used an MHC tetramer based approach to study the differentiation of naïve B cells after exposure to allogeneic antigen (20,21). We found that while anti-donor IgM and IgG was formed after sensitization, few class switched B cells were generated. In addition, a low frequency of antigen-specific B cells formed germinal centers. Together, these data suggest that allogeneic sensitization leads to a relatively low magnitude of B cell differentiation.

We also found that allogeneic sensitization did not result in antigen-specific B cells differentiating into subsets of memory B cells based on the surface receptors CD73, CD80, and PD-L2 (22,23). In contrast, using dimensionality reduction analysis, we found that clusters expressing high levels of CD21 increased after allogeneic sensitization. Using manual gating, we found that the frequency of CD21+ marginal zone phenotype B cells were increased after sensitization. Specifically, the frequency of marginal zone B cells was increased at day 21 and 42 after sensitization.

Marginal zone B cells are defined by their localization in the marginal zone surrounding the B cell follicle of the spleen. The fate decision of transitional B cells to become marginal zone B cells is based on antigen signal strength and dependent on continuous Notch2 signaling (11). Relative to follicular B cells, marginal zone B cells are more sensitive to innate signaling, in particular LPS, and readily differentiate into plasma cells (13–15). Recent reports have implicated marginal zone B cells in the formation of DSA and graft rejection after T cell depletion, adn have shown that Notch2 blockade can prolong cardiac allograft survival. We found that after marginal zone B cell depletion, anti-donor IgG was significantly diminished.

Despite the developmental and functional differences, marginal zone and follicular B cells have overlapping functional capabilities. For example, both populations can form germinal center B cells and ultimately form plasma cells. We found that sensitization of marginal zone depleted mice resulted in fewer germinal center B cells relative to controls. Together, these data suggest a model in which the marginal zone B cells are enriched for allogeneic B cells. Future studies will have to dissect the mechanistic basis for this finding, whether this finding is due to the threshold of activation, localization of antigen delivery, or specificties of antigen-receptors.

## ACKNOWLEDGEMENTS

The authors thank members of the Johns Hopkins Division of Immunology and Vascularized Composite Allograft Laboratory for feedback. This work was supported by NIH AI04616 (S.M.K.).

## Notes

### Competing Interest Statement

The authors have declared no competing interest.

